# Localized prediction of glutamate from whole-brain functional connectivity of the pregenual anterior cingulate cortex

**DOI:** 10.1101/2020.02.26.966259

**Authors:** Louise Martens, Nils B. Kroemer, Vanessa Teckentrup, Lejla Colic, Nicola Palomero-Gallagher, Meng Li, Martin Walter

## Abstract

Local measures of neurotransmitters provide crucial insights into neurobiological changes underlying altered functional connectivity in psychiatric disorders. However, non-invasive neuroimaging techniques such as magnetic resonance spectroscopy (MRS) may cover anatomically and functionally distinct areas, such as *p32* and *p24* of the pregenual anterior cingulate cortex (pgACC). Here, we aimed to overcome this low spatial specificity of MRS by predicting local glutamate and GABA based on functional characteristics and neuroanatomy, using complementary machine learning approaches. Functional connectivity profiles of pgACC area *p32* predicted pgACC glutamate better than chance (R^2^ = .324) and explained more variance compared to area *p24* using both elastic net and partial least squares regression. In contrast, GABA could not be robustly predicted. To summarize, machine learning helps exploit the high resolution of fMRI to improve the interpretation of local neurometabolism. Our augmented multimodal imaging analysis can deliver novel insights into neurobiology by using complementary information.

## 1 Introduction

Since the beginning of the 20^th^ century, the human brain is understood to consist of regions with distinct microarchitecture (Brodmann, 1909; Vogt & Vogt, 1919). Anatomical features, such as cytoarchitecture, myeloarchitecture and receptorarchitecture distinguish cortical areas and highly constrain the local processing capabilities of a region (Cloutman & Lambon Ralph, 2012; Eickhoff et al., 2015; Palomero-Gallagher & Zilles, 2019; Zilles & Palomero-Gallagher, 2017). Cytoarchitecture also shapes a region’s functional repertoire through specific input and output connections, which embed the region in complex distributed networks (Cloutman & Lambon Ralph, 2012). This mesoscopic functional repertoire can be assessed with modern functional magnetic resonance imaging (fMRI) techniques, but methodological and ethical constraints limit us in assessing its relationship to neurometabolism on the microscale *in vivo*. Nevertheless, such linking of functional connectivity (FC) to local processing (e.g. the local excitation/inhibition balance) could provide valuable insights into the pathophysiology of psychiatric disorders.

Though *in vivo* measurements of local metabolism are not possible at a microscopic level, the non-invasive method of proton magnetic resonance spectroscopy (^1^H-MRS) is commonly used to assess local neurotransmitter levels. Available sequences such as PRESS (Bottomley, 1987), STEAM (Frahm, Merboldt, & Hänicke, 1987) and edited sequences such as MEGA-PRESS (Mescher et al., 1998) allow for the detection of glutamate and GABA in single voxels in the human brain. MRS measures of these metabolites may be used to better understand FC changes in psychiatric disorders (Allen et al., 2019; Horn, 2010; Moeller, London, & Northoff, 2016) or pharmacological challenges (Li et al., 2017), but such results may suffer from limited interpretability because of the low spatial specificity of conventional single-voxel spectroscopy. An MRS voxel may cover several known cytoarchitectonically distinct areas (Duncan, Wiebking, & Northoff, 2014). For example, MRS measures of glutamate and GABA follow differential receptor distributions along the anterior cingulate cortex (ACC) (Dou et al., 2013; Li et al., 2017). However, it is not known if the measured concentrations of glutamate and GABA are representative of the entire MRS acquisition voxel and whether the spatial resolution of conventional MRS could be improved by using more fine-grained weights informed by functional imaging.

A region that is considered homogeneous in the MRS literature is the pregenual anterior cingulate cortex (pgACC). This region is part of the default mode network (DMN) of the human brain and has been implicated in the pathophysiology of depression (Salvadore & Singh, 2013). Previous work suggests altered glutamatergic metabolism in the pgACC in patients with major depressive disorder (MDD) (Colic et al., 2019; Horn et al., 2010; Walter et al., 2009) and pgACC levels of a marker of glutamatergic metabolism (glutamate + glutamine, Glx) were correlated with FC between pgACC and insula in patients but not in healthy controls (Horn et al., 2010). Based on its differences in cytoarchitecture and densities of multiple receptors, the pgACC has been divided into six distinct regions: p24a, p24b, pv24c, pd24cd, pd24cv, and p32. These areas were partly merged into a gyral component (areas p24a and p24b into area *p24ab*) and a sulcal component (areas pv24c, pd24cd and pd24cv into area *p24c*) for computation of 3D probabilistic maps (Palomero-Gallagher et al., 2019; Palomero-Gallagher et al., 2008). Studies in non-human primates have shown that homologous areas have distinct structural connectivity patterns (Pandya, Van Hoesen, & Mesulam, 1981). In humans, meta-analytic connectivity modelling showed that these areas have largely distinct functional connectivity patterns, with activation in area *p32* largely associated with tasks involving *Theory of Mind* and cognitive regulation of emotion, and areas *p24ab* and *p24c* with tasks involving the experience of one’s bodily state and action inhibition, respectively (Palomero-Gallagher et al., 2019). In sum, disregarding this heterogeneity during MRS acquisition may hamper the interpretation of links between local metabolism and functional connectivity.

To overcome the problem of low spatial specificity of conventional MRS, we propose a novel, multi-modal approach offering a more nuanced prediction of glutamate and GABA in an MRS voxel. To this end, we employed ROIs originating from a voxel-wise FC-based (‘functional’) parcellation of a pgACC MRS voxel, and a cytoarchitectonic (‘anatomical’) parcellation of the same region (Palomero-Gallagher et al., 2019). We assessed correspondence between the functional and anatomical parcellations and tested whether the prediction of pgACC glutamate or GABA was improved by parcellating the voxel (Fig. 1). Crucially, FC profiles of *p32* but not *p24* could robustly predict local glutamatergic metabolism. We explored why glutamate but not GABA levels were significantly predicted from area FC by examining differential associations between GABAergic and glutamatergic gene co-expression and FC. We then addressed the functional relevance of the differential prediction of pgACC glutamate levels by “decoding” PLSR beta weights. Overall, our results demonstrate that fMRI may improve the spatial specificity of local neurometabolites assessed with MRS.

**Figure 1.**
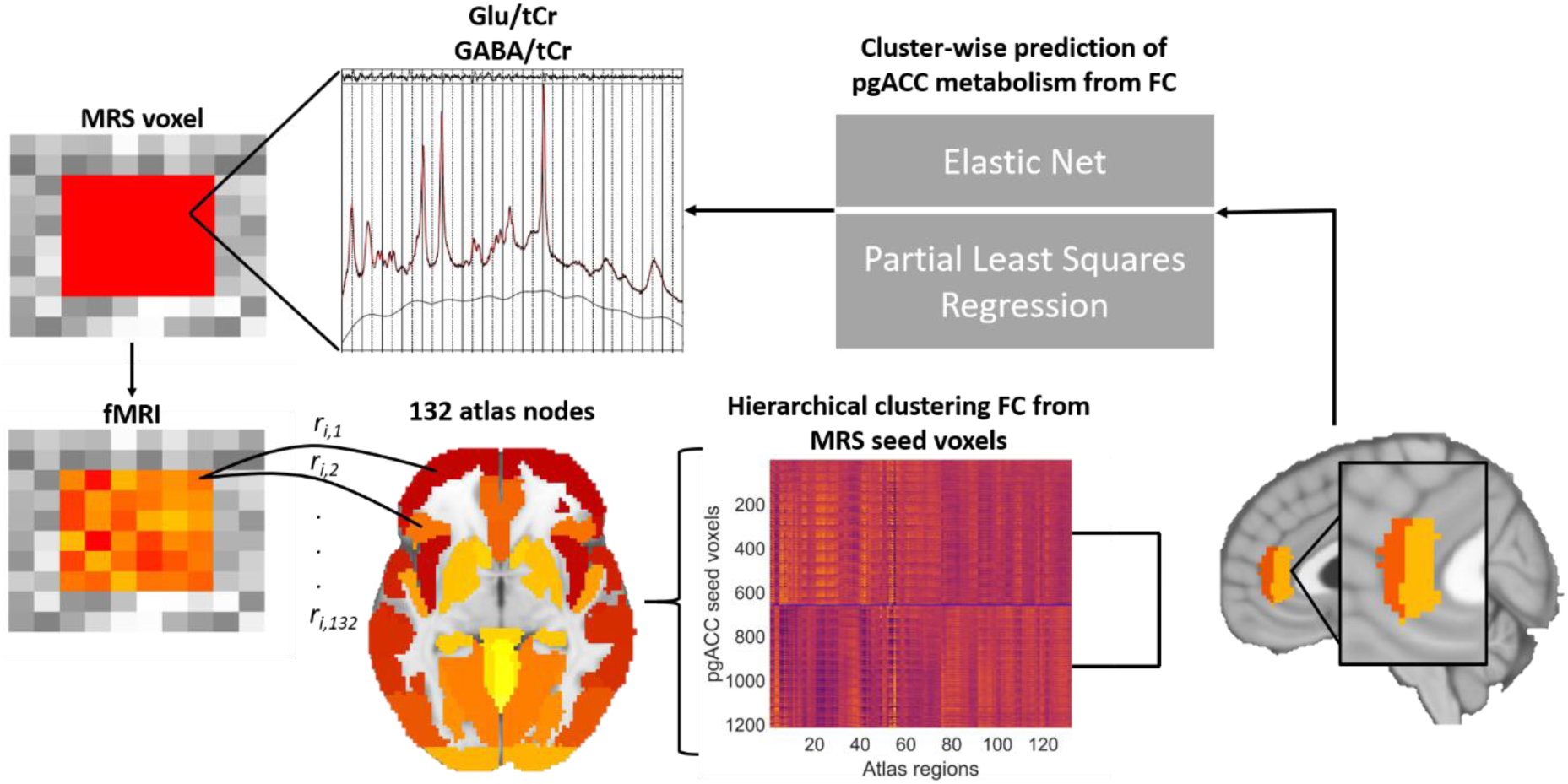
Overview of the primary analysis pipeline. To improve spatial information on local glutamate (Glu) and GABAergic levels measured with MRS, an MRS ROI was parcellated based on seed-based functional connectivity (FC) to 132 atlas nodes. From this hierarchical clustering step, two clusters emerged (Figure S2). Cluster-wise FC profiles were used as input into elastic net and partial least squares regression models to predict GABA/total creatine (tCr) and Glu/tCr residualized for gray matter proportion in participants’ MRS voxels.

## 2. Results

### Functional and anatomical parcellations

For the functional parcellation, we created a group ROI based on participants’ pgACC MRS masks. Participant-specific masks would introduce a bias in the FC profile and could therefore inflate individual associations with neurometabolism. Therefore, we created a composite mask. In addition, we used a recent cytoarchitectonically-informed parcellation of the pgACC as a second, atlas-based ROI parcellation based on 10 post-mortem human samples (Palomero-Gallagher et al., 2019). This anatomical parcellation consists of maximum probability maps (MPMs) of *p24ab*, *p24c*, and *p32*. We parcellated the MRS ROI into two clusters of similar connectivity using hierarchical clustering (for details, see *Materials and Methods* section *Connectivity-based parcellation of the MRS ROI*). Seeds were fMRI voxels within the group MRS ROI; target ROIs were the 132 CONN atlas regions.

We compared the resulting functional clusters to the anatomical parcellation of the pgACC. For this purpose, anatomical maps of *p24* and *p32* were restricted to MRS ROIs. The overlap between the MPMs and the functional clusters was then calculated using Dice coefficients (DC) (cf. Arslan et al., 2018) (Fig. 2c). Cluster 1 overlapped primarily with anatomical area *p32* (DC = .750), but less with area *p24* (DC = .322). Cluster 2 overlapped with anatomical area *p24* (DC = .706) but not with area *p32* (DC = .079). We thus observed good concordance between this region’s parcellation based on local, mesoscale differences and a parcellation based on whole-brain, macroscopic functional differences. Hence, functional clusters 1 and 2 are referred to as functional *p32* and *p24*, respectively, in the remainder of this text.

**Figure 2.**
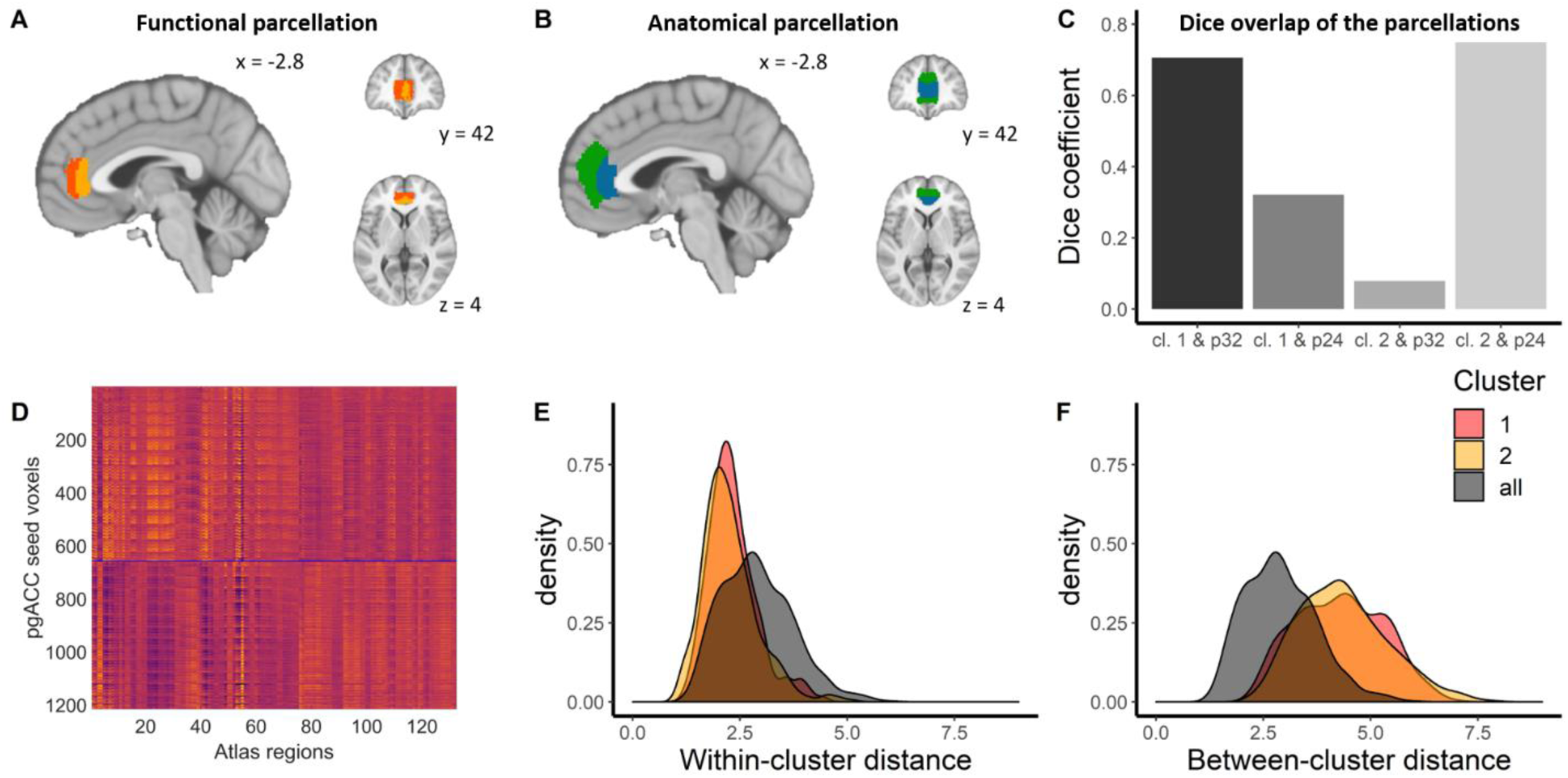
Functional and anatomical parcellations. A: Functional parcellation. Dark orange: cluster 1. Light orange: cluster 2. B: Anatomical parcellation. Green: *p32.* Blue: *p24.* C: Overlap of connectivity-based parcellation results with maximum probability maps from Palomero-Gallagher et al. (2019). Maximum probability maps (MPMs) are masked with the MRS ROI for computation of Dice overlap coefficients. D: Z-scored and de-meaned FC matrix sorted by functional clusters. Dark orange: cluster 1. Light orange: cluster 2. E: Within-cluster distance decreases with functional parcellation (median: cluster 1: 2.25; cluster 2: 2.16; unparcellated: 2.83). F: Between-cluster distance increases with functional parcellation (median: cluster 1: 4.29; cluster 2: 4.27; unparcellated: 2.83).

### Areas covered by the MRS voxel show differential associations with brain networks

To characterize the functional connectivity profiles of each functional area, we calculated area-to-whole-brain connectivity and performed paired t-tests on the results. Fig. 3a-d depicts only those voxels that showed significantly different connectivity (*p* < .05, TFCE). Compared to functional *p24*, functional *p32* showed stronger connectivity to areas that are part of the DMN, including the precuneus and posterior cingulate cortex, inferior parietal lobe, (ventro)medial prefrontal cortex, temporal pole and lateral temporal cortex (Fig. 3a-b). It was also more strongly connected to the inferior frontal gyrus. Functional *p24* had relatively stronger connections to areas that are associated with the ventral attention network (vAt), including the striatum, anterior insula, anterior mid cingulate cortex, and amygdala (Fig. 3c-d). Moreover, according to the t-value distributions of the seed regions, functional *p32* FC is more broadly connected to most Yeo networks (Yeo et al., 2011) compared to functional *p24* FC, which is more specifically associated with the attention networks (vAt and dorsal attention network; dAt) (Fig. 3e).

**Figure 3.**
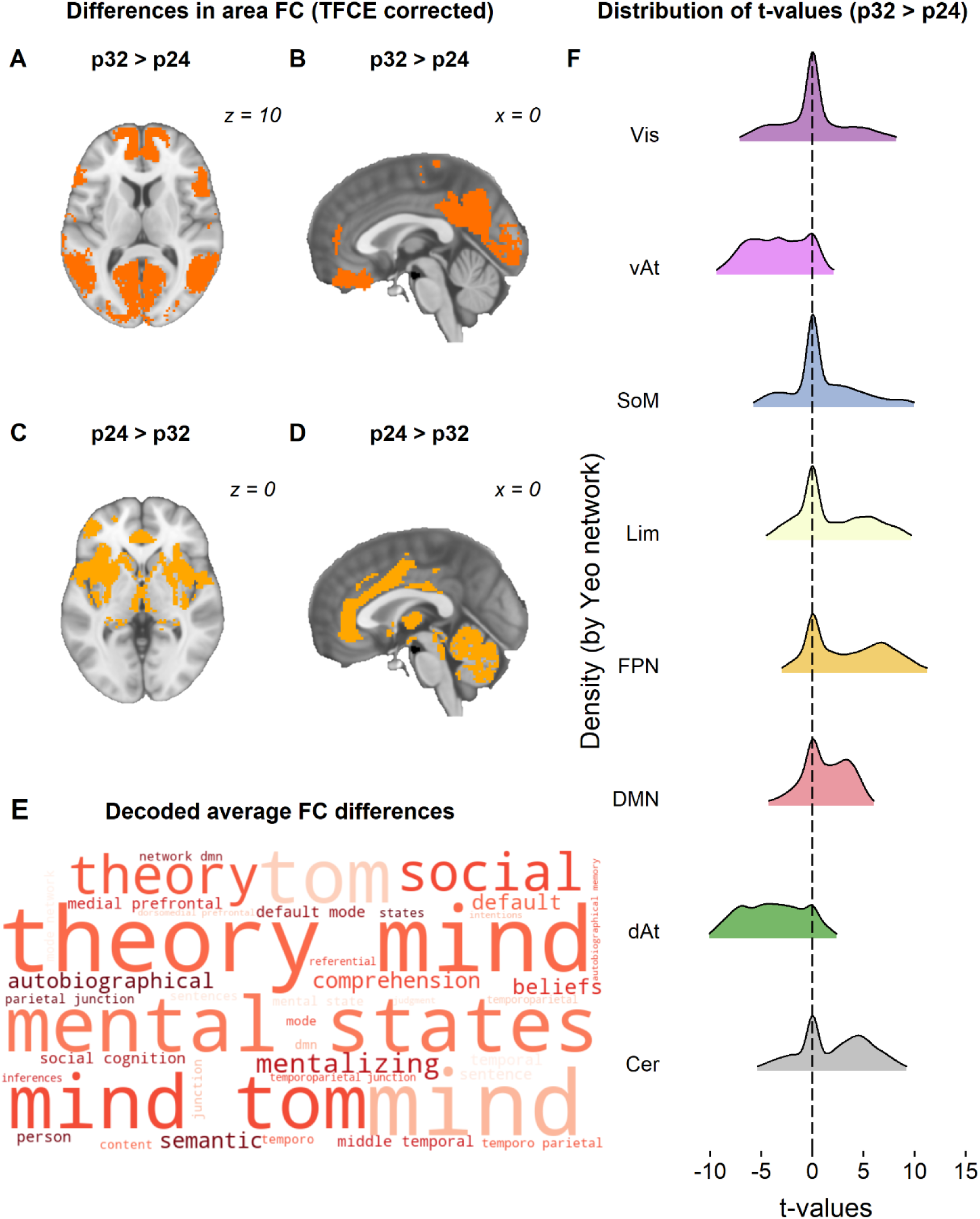
Characterization of FC differences between functional p32 and p24. A-D: Results of paired t-test on whole-brain FC from functional areas, p < .05, TFCE corrected. A & B: functional p32 > functional p24 (orange). C & D: functional p24 > functional p32 (yellow). E: Word cloud of cognitive state terms associated with brain activity in the regions that are more strongly connected to *p32* relative to *p24*. The top 40 terms most strongly associated with this spatial pattern are displayed. F: Density plots of t-values from a paired t-test on whole-brain functional connectivity from functional pgACC areas, p32 > p24, split by Yeo networks and the cerebellar network. Vis = visual network; vAt = ventral attention network; SoM = somatomotor network; Lim = limbic network; FPN = frontoparietal network; DMN = default mode network; dAt = dorsal attention network; Cer = cerebellum.

To further investigate potential associations of average FC differences between functional *p32* and *p24* and cognition, we used the Neurosynth framework (Yarkoni et al., 2011), which comprises neuroimaging data from 14,371 fMRI studies (release 0.7). The decoder toolkit implemented within this framework allows for “decoding” cognitive states from a given (activation) map (Rubin et al., 2017). Compared to *p24, p32* is more strongly connected to a set of regions, that when activated, are associated with cognitive states such as theory of mind, mentalizing, self-referential cognition, and social cognition; cognitive states in which the DMN is thought to be heavily involved (Spreng & Grady, 2010) and that require strong connections to other networks (Barrett & Simmons, 2015; Teckentrup et al., 2019) (Fig. 3f).

### Glutamate is better predicted by *p32* FC compared to *p24* FC

To test whether parcellation of the pgACC MRS ROI into *p32* and *p24* improved the prediction of pgACC glutamate, we employed two complementary machine learning approaches that enforce different degrees of sparsity. Partial least squares regression (PLSR) fits a model based on global information extracted from the feature space and outcome (Zeighami et al., 2017). It is thus able to pick up diffuse, global effects of functional connectivity on local metabolite concentrations. We use elastic net (EN) as a complementary approach to PLSR. EN, in contrast to PLSR, penalizes some regression coefficients (here: FC to target ROIs) to zero, resulting in a sparse model (Pervaiz et al., 2019). EN therefore more strongly enforces spatially specific effects (see *Materials and Methods)*.

First, we used PLSR and EN to test if FC from areas *p24* and *p32* could predict pgACC Glu/tCr (residualized for gray matter proportion) better than expected by chance (Table 1). We found that FC from functional *p32* could be predicted using EN (R^2^ = .324, *p* < .001; Fig. 4b). Although the PLSR model indicated that the FC profile of functional *p32* was associated with pgACC Glu/tCr (R^2^ = .181, *p* = .119; Fig. 4a), this effect did not reach statistical significance. Nevertheless, predicted Glu/tCr values of the EN and PLSR models were highly correlated (R^2^ = .543, *p* < .001), indicating that both methods picked up similar information in the connectivity profiles, and that feature selection in EN was beneficial. Analyses using anatomical *p32* replicated the prediction of Glu by functional parcellation and both models were significantly better than chance (EN: R^2^ = .394, *p* < .001; PLSR: R^2^ = .263, *p* = .023 (Fig. 4c-d)). In contrast to functional and anatomical *p32*, *p24* FC was less consistently predictive of pgACC glutamatergic metabolism. *p24* FC was not predictive of Glu/tCr using PLSR (*p*s > .450) nor using EN (*p*s > .999) (Supplementary Table 1, Fig. 4.). Overall, pgACC glutamatergic metabolism was most reliably predicted from *p32* FC.

**Figure 4.**
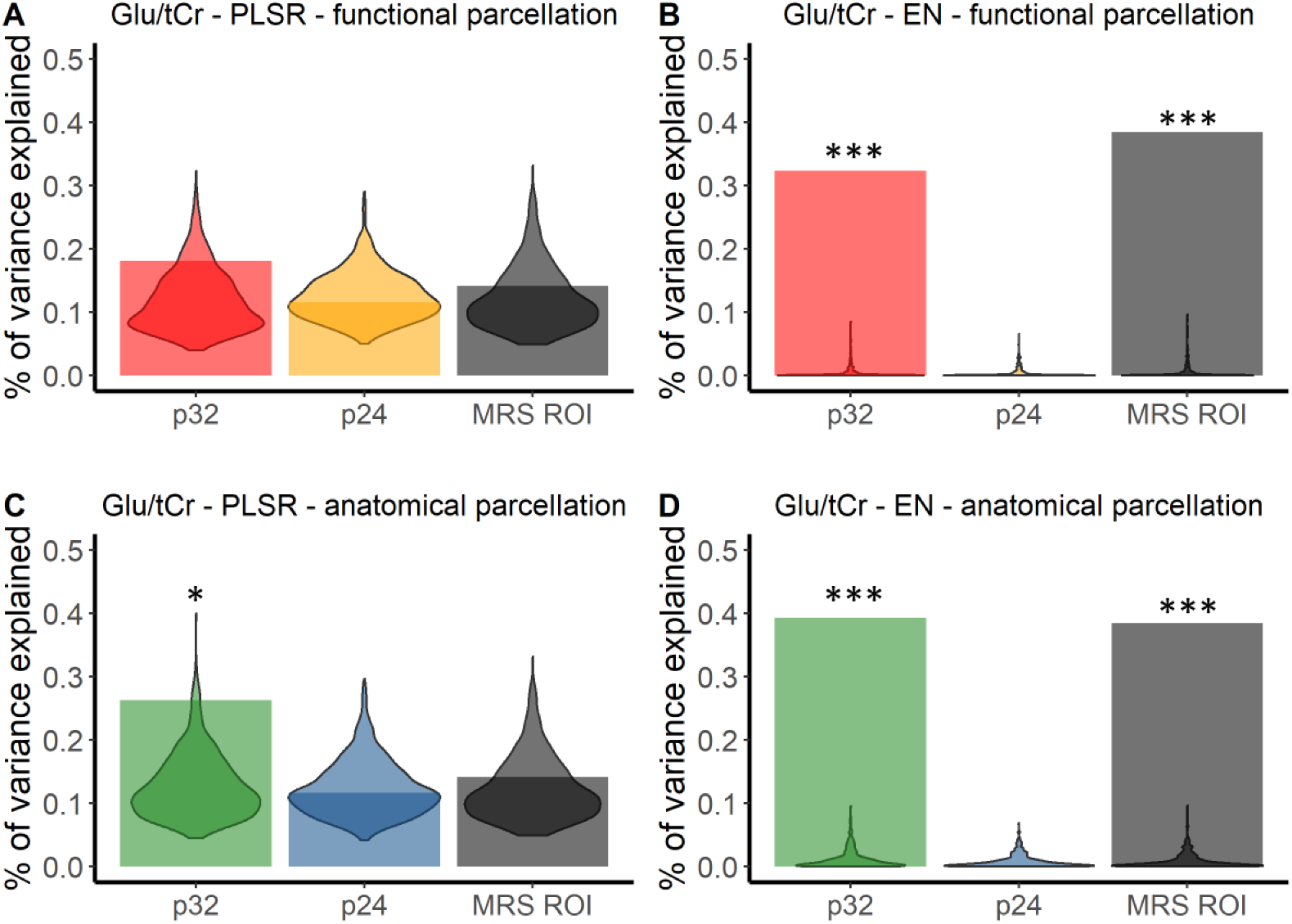
Glutamate levels can be predicted from functional connectivity profiles of *p*32, but not *p*24. Results of partial least squares regression (PLSR) and elastic net (EN) models for Glu/tCr. Violin plots denote the distribution of R^2^ in permutation tests (1000 permutations of metabolite levels). Bars denote R^2^ obtained using Glu/tCr (residualized for gray matter proportion in participants’ MRS voxels) as predictor. * *p* < .05, *** *p* < .001.

FC from the unparcellated MRS ROI could not predict Glu/tCr better than chance using PLSR (R^2^ = .142, *p* = .295), but EN led to comparable results as using *p32* alone (R^2^ = .384, *p* = < .001). To test whether *p32* FC could predict glutamatergic metabolism in the pgACC better than *p24* FC or FC from the unparcellated MRS ROI, we compared the variance explained by two sets of predictors (e.g. *p32* and *p24* FC) using permutation tests. Overall, explained variance of *p32* FC was higher than *p24* FC, demonstrating that *p32* FC was more strongly associated with Glu/tCr compared to *p24* (EN, functional: *p* < .001; PLSR, functional: *p* = .082; EN anatomical: *p* < .001; PLSR anatomical: *p* = .017)*. p32* FC by itself predicted Glu/tCr better or equally well compared to the unparcellated MRS ROI (EN, functional: *p* = .995; PLSR, functional: *p* = .093; EN anatomical: *p* = .110; PLSR anatomical: *p* = .001).

Given the significant correlation between unresidualized Glu/tCr and age in our sample, *r*(86) = −.256, *p* = .016, 95% CI = [−.442, −.049], we explored a possible effect of age on the prediction of Glu/tCr (Supplementary Table 2). Age by itself could be predicted from functional *p32* (R^2^ = .441) and *p24* FC (R^2^ = .310) (*p*s < .001) using EN but not using PLSR (R^2^ = .210, *p* = .066 and R^2^ = .143, *p* = .289, respectively). When the influence of age was accounted for in the EN model by residualizing for age in both FC and metabolite levels, the distinction in predictive ability of the two areas increased. Functional *p32* FC explained numerically more variance in Glu/tCr (R^2^ = .423) compared to the EN model that did not account for age (R^2^ = .403) (Supplementary Table 2).

### pgACC FC is not predictive of pgACC GABA

To test whether pgACC GABAergic neurometabolism could be differentially predicted from FC of parcellated voxels, we repeated the above analyses for GABA/tCr (residualized for voxel gray matter proportion; Supplementary Table 3). A PLSR model built on the FC profile of functional *p24* numerically explained most variance (R2 = 185). However, none of the PLSR or EN models built on FC profiles of anatomical or functional *p24* and *p32* could predict GABA/tCr better than chance. FC from the unparcellated MRS voxel was also not predictive of pgACC GABAergic neurometabolism (*p*s > .05, Fig. 5). Although PLSR models yielded numerically better prediction, none of the functionally or anatomically informed models predicted GABA better than chance.

**Figure 5.**
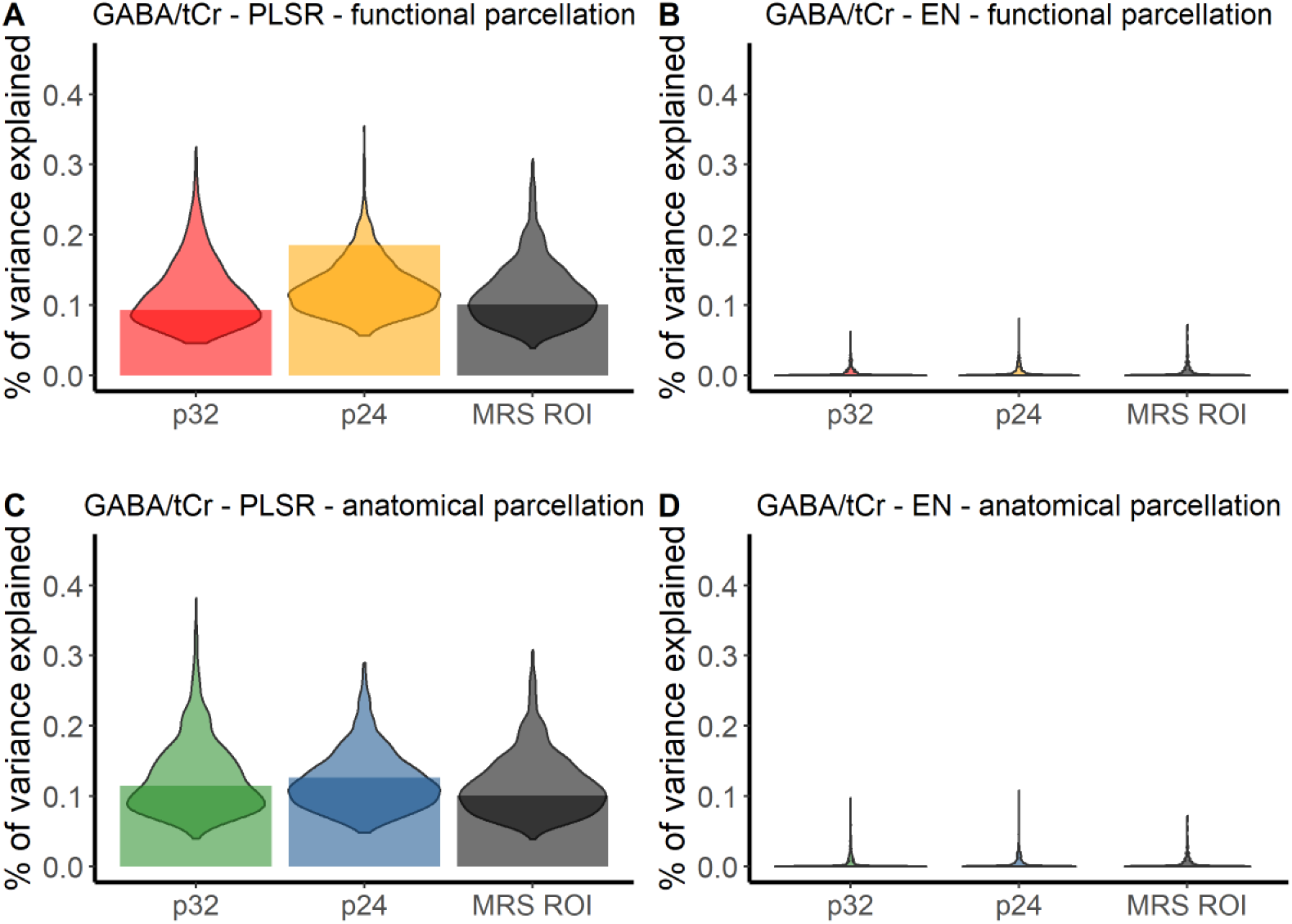
GABA levels cannot be predicted from functional connectivity profiles of the pregenual anterior cingulate cortex. Results of partial least squares regression (PLSR) and elastic net (EN) models for GABA/tCr. Violin plots denote the distribution of R^2^ in permutation tests. Bars denote R^2^ obtained using GABA/tCr as predictor (residualized for gray matter proportion in participants’ MRS voxels).

**Figure 5.**
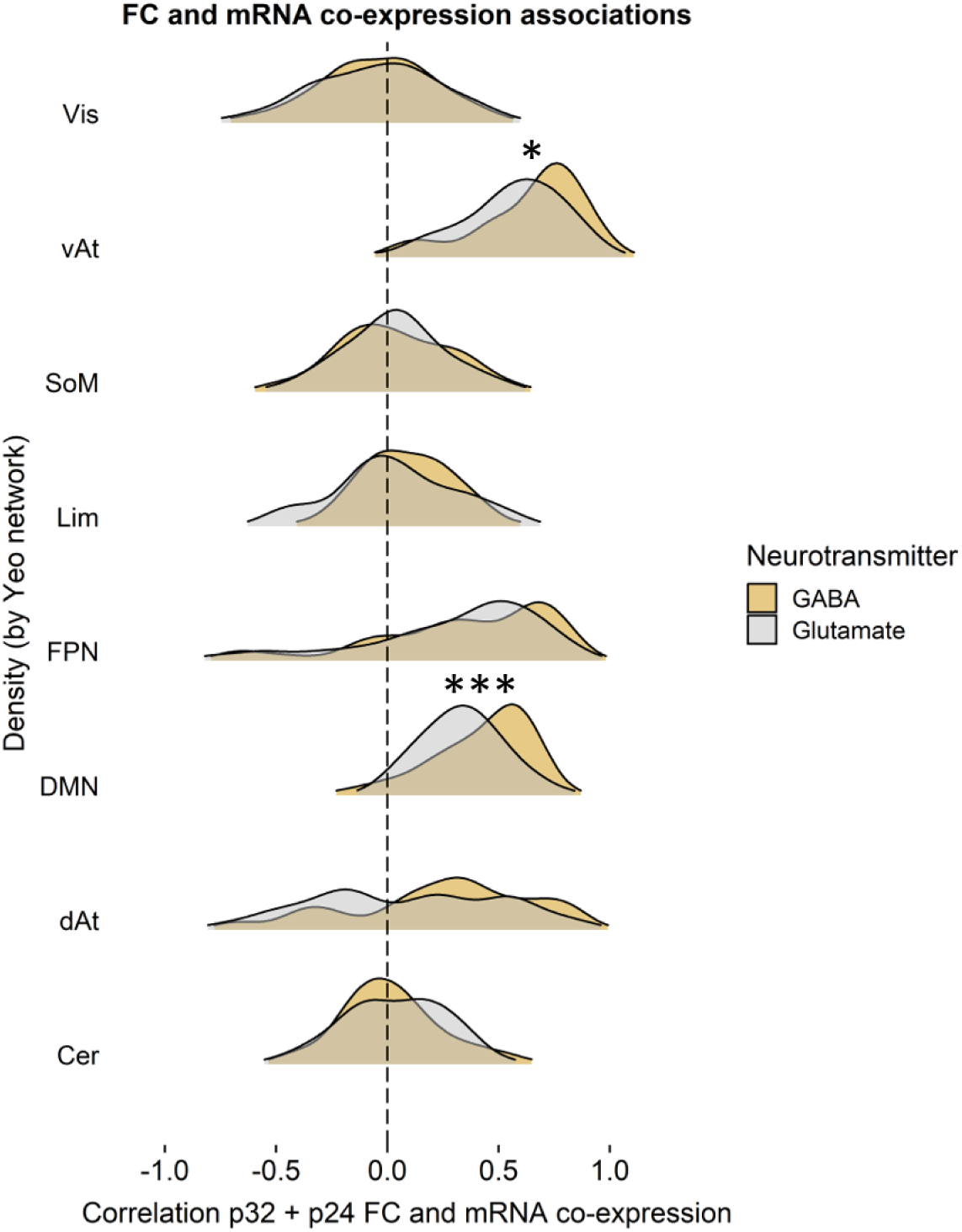
Compared to GABA, there is reduced coupling of glutamate-associated gene co-expression with pregenual anterior cingulate cortex (pgACC) functional connectivity (FC) to vAt and DMN. Density plots show the distribution of glutamate-associated and GABA-associated gene co-expression correlations with *p32* and *p24* FC within the same functional network. Gene co-expression is calculated as the Pearson correlation between pgACC *p24* and *p32* mean mRNA expression with mean mRNA expression in target ROIs. Co-expression-FC correlations are then calculated for each gene and averaged across Yeo networks. * *p* < .05, *** *p* < .001. Vis = visual network; vAt = ventral attention network; SoM = somatomotor network; Lim = limbic network; FPN = frontoparietal network; DMN = default mode network; dAt = dorsal attention network; Cer = cerebellum.

### Co-expression of GABAergic and glutamatergic genes

To investigate why glutamate may be more robustly predicted from area FC than GABA, we explored whether glutamatergic and GABAergic gene co-expression patterns are differentially associated with area FC. Previous work has shown that glutamate and GABA levels across the entire cingulate cortex follow glutamate and GABA receptor fingerprints (Dou et al., 2013). Neurotransmitter receptor density fingerprints shape the local excitatory/inhibitory balance, which influences baseline resting-state functional connectivity (van den Heuvel, Scholtens, & Kahn, 2019). Van den Heuvel et al. (2016) showed an association between the ratio of excitatory and inhibitory gene expression and cortical resting-state FC. In addition, recent work has shown that regions within functional networks share gene expression patterns (‘gene co-expression’) that are distinguishable from those shared within other networks (Anderson et al., 2018; Huntenburg, Bazin, & Margulies, 2018; Richiardi et al., 2015).

The differential prediction of glutamate and GABAergic metabolism may be in part explained by different correlations between resting-state FC patterns of pgACC areas, and canonical glutamatergic and GABAergic gene co-expression patterns obtained from the Allen Human Brain Atlas (see *Materials and Methods* section *Differential mRNA co-expression* for details, and Supplementary Table 4 for the full list of selected genes). We first assessed whether co-expression and pgACC FC were correlated in our sample, as a certain amount of co-expression of GABAergic or glutamatergic genes may be needed for individual variation in metabolism-FC correlations to be meaningful. Intriguingly, two of the networks that were most strongly associated with *p32* and *p24* FC – DMN and vAt – also showed non-zero correlations between gene co-expression for both GABA and glutamate and pgACC FC (Supplementary Table 5). A third network, the frontoparietal network (FPN), also showed non-zero correlations between FC and gene co-expression between pgACC and the network targets. For these three networks, a significant portion of variance in group FC is explained by canonical, structural factors such as gene expression.

As a next step, we investigated differences between GABAergic and glutamatergic co-expression-FC correlations, focusing on the networks for which an association between FC and gene expression was apparent. Co-expression of GABAergic genes within the DMN and vAt was more strongly coupled to FC within those networks relative to glutamate (DMN: D = 0.360, *p* < .001; vAt: D = 0.283, *p* = .011; Supplementary Table 6). This was not the case for the FPN (D = 0.147, *p* = .488). For glutamate-associated genes, this relatively *reduced* coupling between canonical structural and individual functional aspects could leave room for individual variation in Glu levels to influence FC. This could in part explain why compared to GABA, pgACC Glu levels could be better predicted from individuals’ FC profiles.

By comparing FC associations with metabolites versus gene co-expression, we can illustrate this possibility using our data. For each target ROI, the PLSR beta weight represents the strength of the relationship between glutamatergic or GABAergic metabolism and FC to an area. Analogous to gene co-expression (Fig. 5), the distribution of PLSR weights across the vAt was significantly different from zero (Fig. 6A, Supplementary Table 7). For this network, the discrepancy of bootstrapped mean correlations between FC and gene co-expression versus ones between FC and local metabolism was smaller for glutamate (Fig. 6B), such that metabolite-FC correlations were relatively stronger, and co-expression-FC correlations were relatively lower (*p_boot_* = .008).

**Figure 6.**
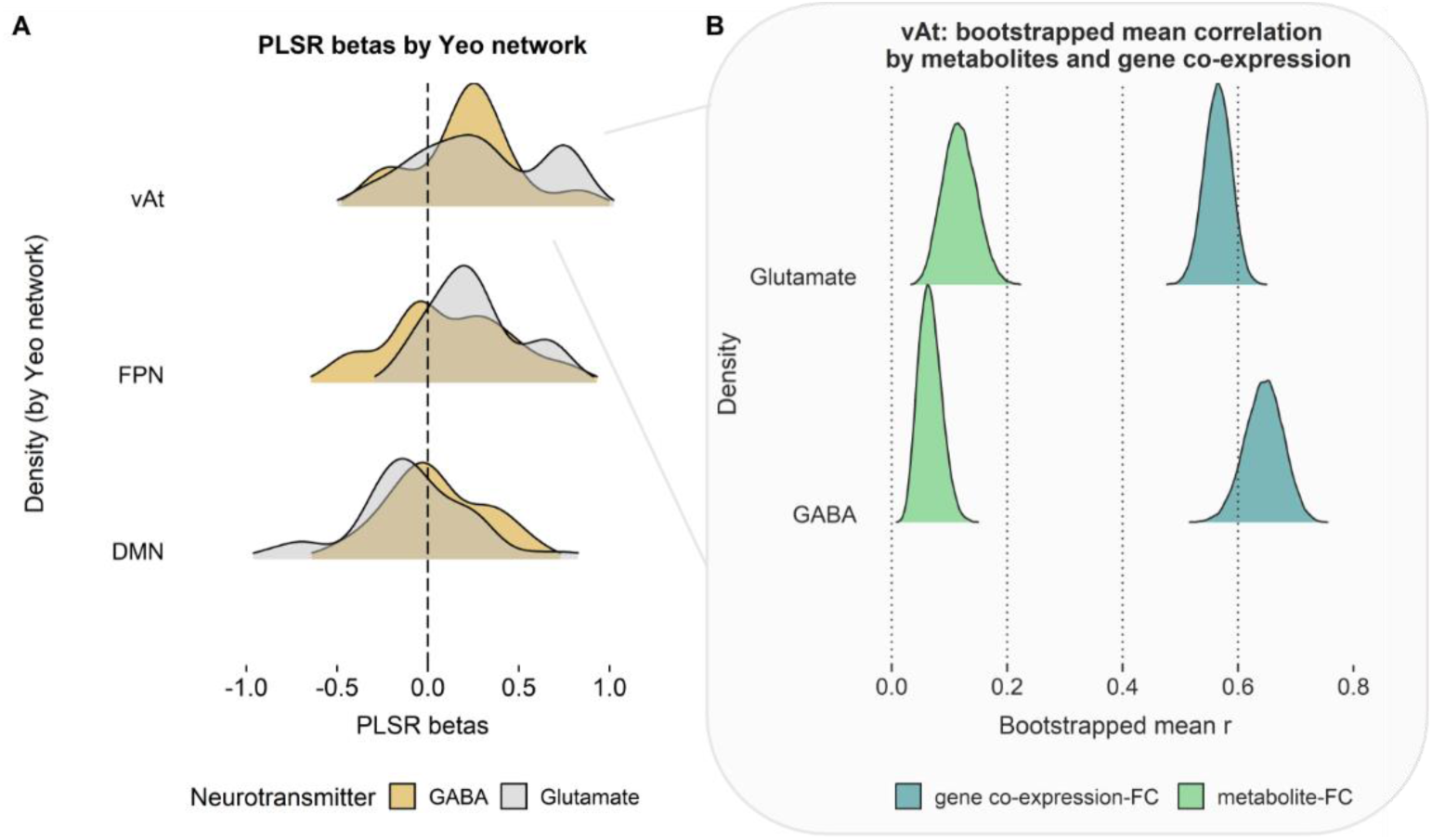
Associations of functional connectivity (FC) profiles with neurometabolites are constrained by canonical gene expression. A. PLSR betas by Yeo network (pooled across functional *p24* and *p32*) show that FC to vAt is associated with GABAergic and especially with glutamatergic metabolism. B. Canonical gene co-expression of GABAergic genes within pgACC and vAt is more strongly correlated to FC than pgACC GABA levels. For Glutamate, this discrepancy is less strong, suggesting that pgACC glutamate levels have more room to influence FC to the vAt. Density plots in (B) display distributions of bootstrapped r values (50,000 iterations). vAt = ventral attention network; FPN = frontoparietal network; DMN = default mode network.

### Cognitive state correlates of *p32* and *p24* FC-glutamate associations

Next, we investigated the potential functional relevance of the differential prediction of pgACC glutamate levels. PLSR beta weight maps for functional *p32* and *p24* show different spatial patterns (Fig. 7A-B, top; see Supplementary Figure 1 for analogous results for anatomical areas). To explore differences between the two pgACC areas, we applied the Neurosynth decoder tool to their respective beta weight maps. Regions for which FC to *p32* is predictive of glutamate, when activated, correspond to cognitive states related to the ventral attention network and cognitive control (Fig. 7A, bottom). In contrast, regions important in the prediction of glutamate from *p24* FC, when activated, reflect cognitive states related to movement and somatosensory experiences (Fig. 7B, bottom). The difference between the two beta maps suggests that relative to *p24*, *p32* FC to the precuneus and posterior cingulate (i.e., DMN regions), anterior insula (vAt) as well as subcortical regions is linearly associated with glutamate levels (Fig. 7C). In general, increased glutamate in the whole pgACC appears to strengthen *p32* FC across a wider functional spectrum compared to *p24*, whose FC coupling is restricted to the somatosensory and motor regions (Fig. 7D).

**Figure 7.**
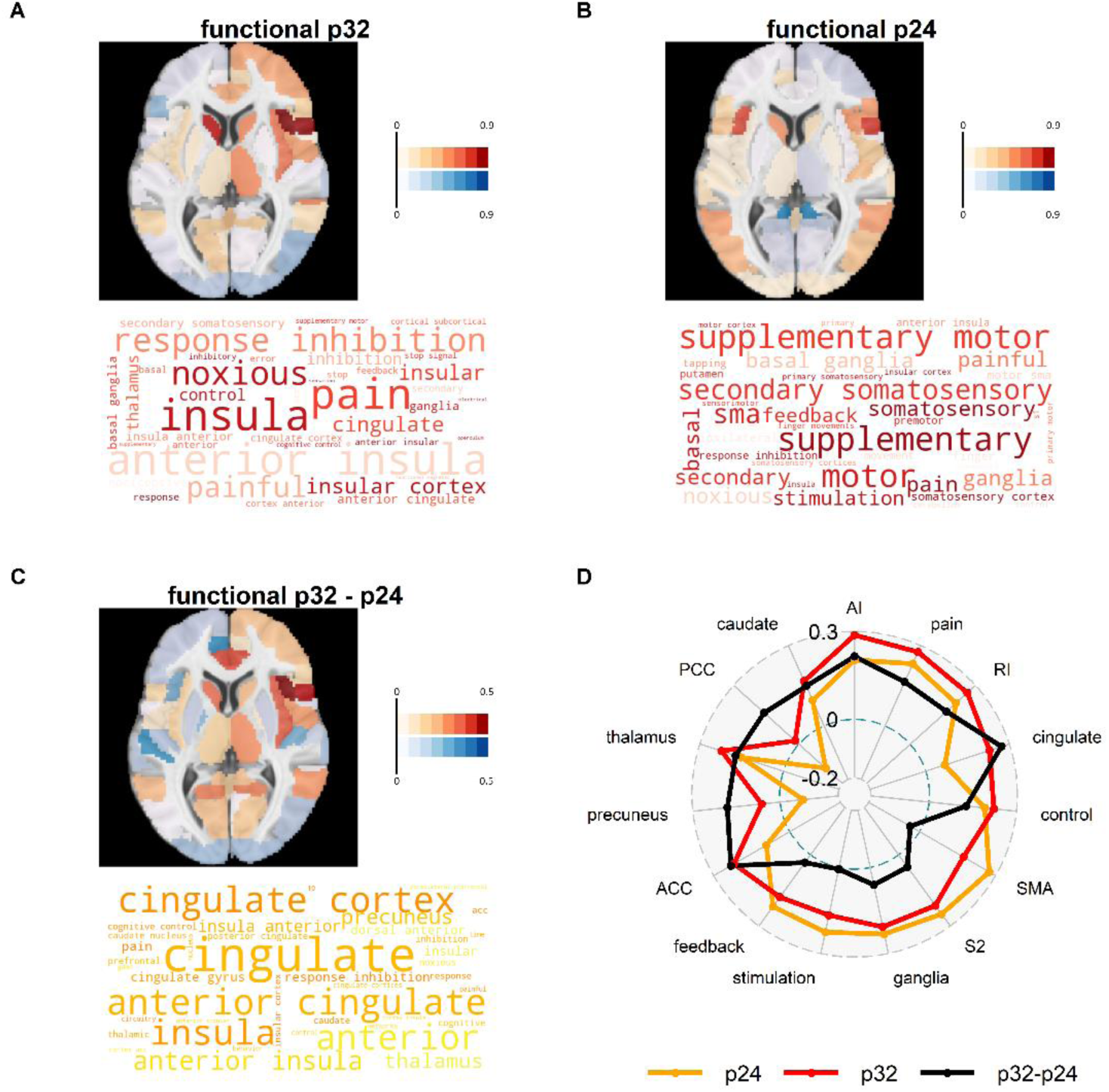
Associations of glutamate loadings with Neurosynth terms show that p32 functional connectivity (FC) extends more to other brain regions. A-C, top: PLSR beta maps show the association between pgACC glutamate levels and pgACC area-to-target FC. A-C, bottom: decoded PLSR beta weight maps show the 40 cognitive state terms most strongly associated with the PLSR beta weight maps. D: spider plot of correlations with the top 5 terms (excluding duplicates) associated with regional brain activity in the spatial pattern represented by PLSR beta weights of *p32*, *p24*, and *p32*-*p24* PLSR, respectively. AI: “anterior insula”; RI: “response inhibition”; SMA: “supplementary motor”; S2: “secondary somatosensory”; ACC: “anterior cingulate”; PCC: “posterior cingulate”.

## 4. Discussion

Psychiatric disorders such as depression and anxiety are characterized by changes in neurometabolites (Colic et al., 2019; Horn et al., 2010; Pollack et al., 2008) and functional connectivity (Mulders et al., 2015). One of the key challenges in biological psychiatry is to understand how changes in neurotransmission lead to changes in brain function so that pharmacological interventions could be used to “normalize” brain function and improve behavioral symptoms (van den Heuvel et al., 2019; Allen et al., 2019; Wang & Krystal, 2014). However, technical limitations of current neuroimaging techniques such as large MRS voxels encompassing heterogeneous areas makes the link of metabolites to function of brain networks non-trivial (a ‘many-to-one mapping problem’, Paulus & Thompson, 2019). Here, we demonstrate that parcellating an MRS region based on functional connectivity or cytoarchitecture improves the prediction of local neurometabolism via global connectivity profiles. Restating the question as a problem of classification rather than localization – can resting-state FC from an area predict local neurometabolism or not? – allowed us to reduce the number of comparisons and address the many-to-one mapping problem. Specifically, we found that area *p32* FC predicts glutamate better than chance irrespective of the parcellation scheme, whereas we did not find converging evidence for the prediction of GABA from FC profiles. Moreover, prediction of glutamate from *p32* FC explained as much or more variance than FC from the unparcellated MRS ROI, while providing additional spatial information. Collectively, our results show that multimodal imaging may help to overcome the fundamental limitations of a single method, as fMRI can improve the spatial specificity of local glutamatergic metabolism assessed with conventional MRS.

Hierarchical clustering of the pgACC MRS voxel recovered clusters in line with anatomical areas *p24* and *p32* (Palomero-Gallagher et al., 2019), with distinct functional connections during rest. The area most predictive of pgACC glutamate, *p32*, is well connected to most functional networks but compared to *p24*, it has particularly strong FC to regions that are part of the DMN. Area *p24* showed relatively stronger connectivity to parts of the ventral attention network or salience network. These findings are in line with previous work in humans (Beckmann, Johansen-Berg, & Rushworth, 2009; Palomero-Gallagher et al., 2019) and non-human primates (Carmichael & Price, 1995; Pandya et al., 1981; Vogt & Pandya, 1987). Moreover, similar distinctions between *p24* and *p32* functional domains were found using direct stimulation of the cortex using stereo-electroencephalography (Caruana et al., 2018). We demonstrated that functional connectivity-based parcellation of an MRS ROI can reflect both cytoarchitectonic areas and well-established connectivity differences.

How should the differential prediction of glutamate FC from the two pgACC areas be interpreted? One possibility is that there is an association between glutamate and p*32* FC, because of the area’s stronger connectivity to the DMN. This intrinsic connectivity network shows the highest metabolic activity at rest (Raichle, 2001). Intriguingly, our findings suggest another possible mechanism. pgACC glutamate increases FC to networks associated with a relatively broad functional range, but particularly to regions of the vAt. Glutamate concentrations in the pgACC may therefore be an important factor contributing to the region’s ability to switch between exteroception (vAt or salience network) and interoception (DMN). The interaction between these networks is often dysregulated in psychiatric disorders (Kaiser et al., 2015; Manoliu et al., 2014; Menon, 2011; Teckentrup et al., 2019). *p32* may therefore be an especially relevant target for future research on metabolic and functional changes in these disorders.

We demonstrated the potential for further applications of this method by prediction of age. Age, unlike gender or voxel gray matter proportion, was highly correlated with glutamate levels. In addition, a wealth of research covers FC changes that occur during aging. Particularly, intrinsic connectivity in the DMN alters with age (Tomasi & Volkow, 2012; Wu et al. 2011; Damoiseaux et al., 2008; Ferreira & Busatto, 2013). Both *p24* FC and *p32* FC were predictive of participants’ age using EN. The successful prediction of age from both areas is encouraging and suggests that this method could be used for differential prediction of other clinical characteristics. It also demonstrates that while *p24* FC is not predictive of glutamate, it is predictive of age. What is more, after regressing out the influence of age from FC and residualized Glu/tCr, the variance explained in metabolism by *p24* was reduced, whereas variance explained by *p32* increased compared to the model in which age was not accounted for.

In contrast to the considerable predictive power for glutamate, we did not find converging evidence for the prediction of local GABAergic metabolism via global connectivity profiles. Previous work demonstrated a positive correlation between GABA levels in the pgACC and negative BOLD responses upon emotional stimulus presentation (Northoff et al., 2007), suggesting a link between GABA and the potential to downregulate DMN activity with increasing cognitive load. As our measurements were acquired at rest, it may be that an association between GABA and FC becomes apparent after stronger recruitment of task-positive networks. Another explanation relates to the complex association between GABAergic metabolism and the BOLD response. Depending on the brain region and network, an increase in inhibitory activity may or may not lead to increased BOLD response (Bartels, Logothetis, & Moutoussis, 2008; Logothetis, 2008). Our results showed that GABA-associated genes were more tightly linked to pgACC FC compared to glutamate-associated genes. GABA co-expression-FC relationships may thus be less variable across individuals. GABA modulates glutamatergic excitation by acting on pyramidal neurons in cortical microcircuits and local GABA concentrations may therefore represent mostly local processes (Isaacson & Scanziani, 2011; Logothetis, 2008; Buzsáki, Kaila, & Raichle, 2007). To summarize, our results suggest that pgACC measures of GABA are unlikely to be associated with patterns of long-range functional connectivity at rest, calling for alternative techniques in the future.

Strikingly, while functional and anatomical parcellation performed similarly, there were vast differences in performance between EN and PLSR models. For the region most predictive of glutamate, *p32*, predicted values from EN and PLSR were highly correlated, suggesting they pick up similar information. Nevertheless, EN models provide a more conclusive answer on whether a metabolite can be predicted or not. In case there is no relationship between outcome and predictors, for EN, the built-in 10-fold cross-validation will lead to models with all predictors reduced to zero, because there is no lambda yielding a better than chance out-of-fold prediction. In such situations, PLSR models can still result in high explained variance, because these are not equipped with a way to penalize predictors that do not remain predictive in held-out folds. EN models have previously been used to predict behavior (Kashyap et al., 2019) and disease (Teipel et al., 2017) from neuroimaging data. This method outperforms multiple regression (Jollans et al., 2019; Teipel et al., 2017) and also frequently outperforms other machine learning techniques in cases where the number of participants is similar or smaller than the number of datapoints (Jollans et al., 2019). Overall, based on our results, combining MRS measures of local neurometabolites with resting-state FC might be promising to identify candidate regions or networks.

The results presented here must be considered in light of several limitations. First, the mean Dice overlap between MRS voxels and the functional areas was significantly greater for *p24*. Nevertheless, functional area *p24* was *less* strongly associated with glutamatergic metabolism compared to functional area *p32*. In addition, there was no significant relationship between participants’ MRS voxel overlap with functional clusters and their glutamate or GABA levels. Therefore, it is unlikely that the larger MRS voxel overlap with cluster *p24* had a significant influence on our results. Second, as in most MRS studies, there are potential confounds in the quantification of neurometabolites. While GABA is more challenging to reliably quantify compared to glutamate, it is unlikely that data quality played a role in our findings. At high field strengths like 7 Tesla, increased signal dispersion allows for the separation of GABA peaks from larger, overlapping resonances. Moreover, after our stringent quality control, only five participants had to be excluded from the GABA analyses. With regard to glutamate, MRS measures are not limited to glutamate as neurotransmitter. Glutamate fulfills other, metabolic roles in the cell, including protein synthesis and energy metabolism, which cannot be separated from as neurotransmitter (Rae, 2014). It also appears that vesicular glutamate is not detectable by MRS (Kauppinen and Williams, 1991). The generalizability of the models needs to be assessed in an independent dataset. For this purpose, beta maps in MNI space are available for download: https://neurovault.org/collections/CWXNJGOY/. Last, the gene expression data set was obtained from an independent sample (Allen Human Brain Atlas) consisting of six donor brains. At present, it is unclear whether gene expression and MRS measures are well aligned and whether gene co-expression generalizes to a wider population of healthy living adults. Notwithstanding, our study shows that associations of FC with canonical gene expression can provide crucial insights into neurobiology.

To summarize, we demonstrated that combining complementary information from different neuroimaging modalities (MRS and fMRI) can provide incremental spatial information on the relationship between function and neurometabolism, by capitalizing on the higher resolution of fMRI. Our results show that *p32* is more predictive of pgACC glutamate compared to *p24* and suggest that although *p32* as a DMN node is strongly connected to most networks, pgACC glutamate concentrations are particularly associated with *p32* FC to the ventral attention or salience network. Unlike glutamate, GABA could not be reliably predicted from pgACC FC, as canonical GABAergic co-expression may be more influential. As smaller voxel sizes reduce SNR, this approach could be used as an alternative to extract more localized information about key neurometabolites and can be particularly informative when the MRS ROI cannot be restricted to one functionally distinct area. Importantly, this method could be applied to other multimodal datasets, including EEG-fMRI or PET-fMRI to improve the spatial resolution of inferences. Crucially, our novel combination of techniques can be readily used in existing datasets to uncover more spatially specific relationships between functional connectivity underlying neurometabolism in health and disease. Thus, a broader application of interpretable machine learning methods may lead to a better understanding of neurobiological mechanisms of common psychiatric disorders.

## Materials and Methods

### Participants

We included 143 healthy participants in this study. Data were pooled from three studies. All participants were screened for prior and current neurological or psychiatric illness using the German version 5.0.0 of the M.I.N.I. Mini International Neuropsychiatric Interview (Ackenheil et al., 1999). All study procedures were approved by the ethical committee of the University of Magdeburg and conformed with the Declaration of Helsinki.

Several participants participated in more than one MRS acquisition. In these cases, we discarded the measurement that did not meet initial quality criteria or, in case multiple measurements of the same participant were of sufficient quality, we selected the measurement with the best MRS and/or fMRI data quality in advance of the current analysis.

The final sample consisted of 88 participants with good quality resting-state and MRS data (age = 28.81 ± 9.02; 35 females) for analysis of Glu/tCr. For GABA/tCr, one female and four male participants could not be included in the analyses because of CRLBs exceeding the cut-off of 20, leading to a reduced sample size of 84 participants.

### Data acquisition and preprocessing

#### MRS

Ultra-high field data were acquired on a 7T MAGNETOM scanner equipped with a 32-channel head coil (Siemens, Erlangen, Germany). Before MRS measurements, an MPRAGE T1-weighted scan was acquired. The echo time (TE) was 2.73 ms, repetition time (TR) was 2300 ms, and inversion time (TI) was 1050 ms. In-plane field of view (FOV) was 256 mm. The flip angle was set to 5°. Images were acquired with a bandwidth of 150 Hz/pixel and 0.8 mm isotropic image resolution.

Participants’ T1-weighted scans were used for accurate placement of the pgACC voxel, according to an established protocol of anatomical landmarks described in Dou et al. (2013). Briefly, the pgACC voxel (10 x 20 x 20 mm^3^) was placed in the bilateral pgACC and centered on the sagittal midline to ensure maximal coverage of cingulate gray matter. Automatic shim routines were used to optimize B_0_ homogeneity. We applied a stimulated-echo acquisition mode (STEAM) sequence with variable-rate selective excitation (VERSE) RF pulse (Dou et al., 2013) with short TE/mixing time (TM) (20 ms/10 ms) and TR = 3000 ms. Metabolite spectra were acquired with 128 averages. A single water reference signal was acquired for eddy current correction. The bandwidth was 2800 Hz and the acquisition time for one image was 731 ms.

MRS data were fitted in LCModel V6.3.0 (Stephen Provencher, Oakville, ON, Canada; Provencher, 2001). The basis set used for fitting included Creatine, Glutamate, Myo-Inositol, Lactate, N-acetylaspartate, Phosphocholine, Taurine, Aspartate, γ-Aminobutyric acid (GABA), Glutamine, Glucose, Alanine, N-acetylaspartyl-glutamate, Phosphocreatine, Scyllo-inositol, Acetate, Succinate, Phosphorylethanolamine, Glutathione, Citrate, and Glycerophosphocholine. In all analyses, we used Glutamate and GABA as a ratio to total Creatine, i.e. Creatine + Phosphocreatine (tCr). Metabolite values were considered of insufficient quality if signal-to-noise (SNR) was smaller than 20, if linewidth (full-width-at-half-maximum) was larger than 24 Hz or if the CRLB was smaller than 20%.

To account for differences in voxel gray matter (GM) content, we segmented participants’ T1-weighted images using VBM in the CAT12 toolbox for SPM (http://dbm.neuro.uni-jena.de/cat/index.html#VBM, Ashburner & Friston, 2005). MRS voxels were normalized to MNI space using the forward deformation field produced during segmentation and normalization of the structural scans. Participants’ normalized gray matter tissue probability map produced by VBM were then masked with their normalized MRS mask, and the percentage of probable voxel GM content was calculated. We regressed out voxel GM content from Glu/tCr and GABA/tCr values and performed all further analyses on the residuals.

#### rs-fMRI acquisition

For the acquisition of resting-state fMRI data, participants were instructed to keep their eyes closed and think of nothing in particular. Acquisition time for this measurement was 13:18 min and TR/TE were 2800 ms and 22 ms, respectively. The image resolution was 2 mm isotropic and the in-plane FOV was 212 mm. The flip angle was 80°. Sixty-two slices were acquired for a total of 280 volumes. In-plane parallel imaging was done with GRAPPA image reconstruction (Griswold et al., 2002) acceleration factor 3. The first 10 volumes of resting-state data were discarded to reach steady state.

#### ROI definition

For functional parcellation, we created an ROI based on participants’ pgACC MRS masks. Participant-specific masks would introduce a bias in the functional connectivity profile and could therefore inflate associations with local neurometabolism. For this reason, we created a composite mask. We resampled normalized participants’ MRS masks to the space and resolution of functional images (2 x 2 x 2 mm^3^). From the resulting masks, we created a composite group MRS ROI such that each voxel within the ROI was contained in the normalized MRS mask of at least two participants (i.e., threshold for inclusion of a voxel: > 1). In addition, we used a recently published anatomical parcellation of the pgACC as a second, atlas-based ROI parcellation (Palomero-Gallagher et al., 2019). This parcellation consists of maximum probability maps of areas *p24ab*, *p24c*, and *p32*. The delineation of the areas reflects cytoarchitectonic differences and is based on 10 post-mortem human samples.

#### rs-fMRI pre-processing

Pre-processing of resting-state data was done using the CONN toolbox (Whitfield-Gabrieli & Nieto-Castanon, 2012). Briefly, images were realigned and unwarped (motion correction) and slice-time corrected. Functional and structural images were then segmented using default tissue probability maps and normalized to MNI using direct normalization and resampled to 2 mm isotropic voxel size. To take the fullest advantage of the gained spatial resolution as a result of using ultra-high field strength, we did not apply spatial smoothing. This approach also allowed us to limit the calculation of seed-based FC to the MRS ROI. Further denoising was performed using custom MATLAB scripts. In this step, time series from voxels with either a GM probability of .35 or higher, or from those voxels falling within the MRS ROI, were z-scored, despiked, and subjected to quadratic detrending, after which six motion parameters (estimated during the realignment step) and mean white matter signal were regressed out. We did not perform bandpass-filtering of the time series, as most high-frequency fluctuations related to physiological noise were likely to have been removed by regressing out the mean white matter signal (Kahnt et al., 2012) and low-frequency drifts were removed in the detrending step (Tanabe et al., 2002).

#### FC calculation

FC was calculated separately for the group MRS ROI and the anatomical ROI. As seed voxels, we selected only those fMRI voxels in ROIs. To be able to interpret the resulting cluster solution in anatomically informed ways and to reduce the feature space to *k* closer to our N (88), we selected the 132 CONN atlas nodes as target ROIs. These include Harvard-Oxford cortical and subcortical ROIs (Desikan et al., 2006) as well as cerebellar ROIs from the AAL atlas (Tzourio-Mazoyer et al., 2002). Functional connectivity between seed time series and mean target time series was calculated as the Pearson correlation between the two. This resulted in a three-dimensional connectivity matrix with 1216 (MRS ROI) or 2160 (atlas-based ROI) rows (seeds), 132 columns (target ROIs), and 88 participants in the z-dimension.

### Connectivity-based parcellation of the MRS ROI

We parcellated the pgACC MRS ROI into clusters of similar connectivity using resting-state functional connectivity-based parcellation. The aim of this method is to decrease within-cluster distance, and to increase between-cluster distance (Eickhoff et al., 2015). First, the seed-by-ROI-by-participant matrix was Fisher z-transformed and averaged across participants (cf. Kahnt et al., 2012). We then computed the correlation between every seed’s connectivity profiles. To finally parcellate contained voxels, we created a similarity matrix (*N_seed_* x *N_seed_*) with Pearson correlation coefficient as the distance measure.

To assess the functional hierarchy within the MRS seed voxel, the resulting similarity matrix was then subjected to hierarchical clustering to cluster functional voxels according to their similarity in terms of whole-brain functional connectivity (Johnson, 1967). A major advantage of hierarchical clustering is that unlike, for example, the popular *k*-means algorithm, a hierarchical approach does not require a predefined number of clusters. The dendrogram may be cut at any level, with *k* + 1 clusters always nested in *k* clusters (Cloutman & Lambon Ralph, 2012; Eickhoff et al., 2015). We used the “average distance” linkage algorithm for clustering. Based on inspection of the dendrogram and the FC matrices, a two-cluster solution was found to be optimal for the overarching goal of the study. Note that due to the hierarchical clustering algorithm, more fine-grained parcellations could be tested to further localize predictive voxel clusters within their cluster branch. Yet, this would come at the cost of multiple-comparison correction and makes specificity incrementally harder to demonstrate. Thus, we decided to focus solely on the two-cluster solution to demonstrate the utility of the method in principle.

On account of their functional similarity, we averaged the functional connectivity to target regions across all seed voxels for each cluster, to reduce the number of features in subsequent analyses. This resulted in separate matrices for each cluster, representing the cluster-to-target FC for each participant. To assess effectivity of functional parcellation we calculated within-cluster distance (sum of squares) and compared it to between-cluster distance (sum of squares). Both within- and between-cluster distance were compared to whole MRS ROI distance to mean FC.

### Cluster validity

Studies using tractographic and functional connectivity-based parcellation showed substantial correspondence between their parcellation and probabilistic cytoarchitectonic maps (Blumensath et al., 2013; Gordon et al., 2016; Wig, Laumann, & Petersen, 2014). Others have provided qualitative evidence (visual inspection) for overlap between cytoarchitecture and connectivity-based-derived parcellation (e.g. Balsters, Mantini, & Wenderoth, 2018; Beckmann, Johansen-Berg, & Rushworth, 2009; Gordon et al., 2016).

We compared our functional clusters to the anatomical parcellation of the pgACC. We summed the maximum probability maps (MPMs) of clusters p24ab and p24c to create a mask of area p24. The MPMs of areas p24 and p32 were then resampled to the space of our functional clusters. MPMs were restricted to MRS ROIs. The overlap between the MPMs and the functional clusters was then calculated using Dice coefficients (DC) (cf. Arslan et al., 2018).

### Influence of individual participants’ voxel placement

To investigate a possible mediating influence of participants’ exact voxel placement on neurometabolite prediction, we compared the means of overlap of participants’ individual MRS masks with functional *p32* (M_DC_ = .357, SD_DC_ = .125) and with functional *p24* (M_DC_ = .482, SD_DC_ = .112), *t*(87) = −5.108, *p* < .001. The Dice overlap between individual MRS voxels and functional *p32* did not correlate with local Glu/tCr (r(86) = −0.952, *p* = .344) or GABA/tCr (r(81) = 1.311, *p* = .193), confirming that individual participants’ voxel placement did not influence our results.

### Statistical analyses

Correlational analyses and t-tests were performed in R (Version 3.5.0) with the RStudio IDE (Version 1.0.136). All other statistical analyses were performed in MATLAB 2019a. α was set to .05. Because the methods - elastic net and partial least squares regression - are not independent, but complementary, we did not correct for multiple comparisons.

#### Demographics

To assess the influence of possible confounders, for both Glu/tCr and GABA/tCr, we calculated Pearson’s correlations with age and gray matter proportions. To test for a difference between Glu/tCr and GABA/tCr between male and female participants, we performed a Welch Two Sample t-test.

#### Characterization of functional ROI FC profiles

We aimed to characterize the functional connectivity profiles of the functional areas *p32* and *p24*. To this end, we calculated participant-wise mean time series for each area, and calculated Pearson correlations with participant-wise whole-brain time series. We performed a paired t-test on both areas to test which brain areas were significantly more functionally connected to either functional area, compared to the other functional area. This analysis was conducted using an implementation of threshold free cluster enhancement (TFCE) in MATLAB (https://github.com/markallenthornton/MatlabTFCE). The threshold for significance was set at .05, TFCE-corrected. For visualization purposes, we repeated the paired t-test in MATLAB, and plotted the unthresholded t-values (*p32* > *p24*) by Yeo network (Fig. 3e).

#### Neurosynth decoding

To further illustrate the difference between functional *p32* and *p24* FC, we “decoded” average FC difference maps (*p32* – *p24*) using data from Neurosynth framework (Yarkoni et al., 2011). The Neurosynth framework comprises neuroimaging data and text extracted from 14,371 fMRI studies. The decoder toolkit implemented within this framework allows for “decoding” cognitive states from a given (activation) map (Rubin et al., 2017). Using this toolkit, we correlated the average FC difference map with all cognitive state maps available in the Neurosynth database (release 0.7). Each voxel in a cognitive state posterior probability map reflects the likelihood that a cognitive state term is used in a study if the voxel is activated (Quintana et al., 2019; Rubin et al., 2017). The top 40 terms correlated with the difference map are displayed in a word cloud (Fig. 3f).

#### Partial-least squares regression

To predict Glu/tCr and GABA/tCr from functional and anatomical *p32* and *p24* FC, we employed partial least squares regression (PLSR) (Krishnan et al., 2011; McIntosh & Lobaugh, 2004). PLSR is a method that is particularly suitable for high-dimensional regression problems, where the number of parameters is larger than the number of samples. PLSR projects the predictor variables into a latent space, while optimizing the prediction of the outcome. PLSR is similar to *principal component regression* (PCR), but while PCR only constructs components based on the captured variance of the predictors, PLSR aims to maximize covariance between factors extracted from predictors and the outcome variable.

PLSR was run in MATLAB using the *plsregress* function, which uses the SIMPLS algorithm (De Jong, 1993). New PLSR models were constructed for each area and each metabolite. One component was retained in each analysis. As outcome variables we used the residuals from a regression model where voxel GM proportion predicted either Glu/tCr or GABA/tCr. The resulting residuals were subsequently z-scored. Predictors were the *p24* or *p32* FC values for each participant. To statistically assess the obtained model fit (residual sum of squares, Abdi, 2010), we performed permutation tests with 1000 permutations of the outcome measure. To test whether one region explained more variance than the other area or the unparcellated MRS ROI, we first calculated the difference in R^2^ between two models (e.g. *p24* and *p32*). Then, we created a null distribution by running EN models with 1000 permutations for both ROIs simultaneously, using the same permuted outcome vector for both ROIs. We assessed statistical significance by comparing the true difference in R^2^ to the permutation distribution.

#### Elastic net

Elastic net (EN) models are a combination of least-absolute-shrinkage-and-selection-operator (LASSO) regression and ridge regression. EN models perform variable selection while simultaneously shrinking regression coefficients to prevent overfitting. We use EN as a complementary approach to PLSR. PLSR fits a model based on global information extracted from the feature space and outcome. It is thus able to pick up diffuse, global effects of functional connectivity on local metabolite concentrations. EN, in contrast, penalizes some regression coefficients (here: FC to target ROIs) to zero, resulting in a sparse model. EN therefore more strongly enforces spatially specific effects.

Residualized GABA/tCr and residualized Glu/tCr were predicted using EN models fit for both functional and anatomical *p24* and *p32* ROIs. Alpha, the weighing term of LASSO and ridge regression in the EN, was set *a priori* to 0.5. The mean squared error of the model fit was estimated using 10-fold cross-validation. Lambda was set to the value with minimum cross-validation error. Robust beta weights used for predicting metabolite concentrations were derived from the median of 20 EN iterations. Descriptive statistics of model fit are given by the R^2^ of predicted metabolite values and actual metabolite values. Analogous to the PLSR models, we assessed the model fit (R^2^) using a permutation test where the order of the outcome vector was randomly permuted (N = 1000). Also analogous to the PLSR models, we compared the difference in explained variance between two areas using permutation tests with 1000 permutations of the outcome vector. Given the significant association between Glu/tCr and age in our sample, we ran EN and PLSR models to predict age rather than glutamatergic metabolism and compared model performance to models where age was regressed out of Glu/tCr and the predictors.

#### Differential mRNA co-expression

To explore what could explain the differential association between glutamatergic metabolism and FC from the two functional areas, we investigated gene co-expression. mRNA expression data from six donor brains were obtained from the Allen Human Brain Atlas (httpt://human.brain-map.org). We selected a subset of genes that are associated with glutamatergic and GABAergic transmission, receptors, transport, and metabolism (see Supplementary Table 4 for the full list of selected genes). When multiple probes were available for the same mRNA, we selected the probe with the highest differential stability, i.e. the probe with the lowest spatial variability between donors (Hawrylycz et al., 2015; Quintana et al., 2019).

We investigated correlations between functional connectivity and gene co-expression within networks. Previous research has demonstrated that regions that are functionally connected (i.e. are part of functional, distributed networks) show similar gene expression patterns distinct from that shared within other networks (Huntenburg et al., 2018; Richiardi et al., 2015; Anderson et al., 2018). The different resting-state functional connectivity from the two areas to the rest of the brain may be in part be explained by glutamatergic or GABAergic gene co-expression patterns. Therefore, we further explored whether gene co-expression between the pgACC and the CONN atlas regions is differentially correlated with functional connectivity from functional p32 and p24.

To this end, we used the gene set described above and computed mRNA expression maps according to Quintana et al. (2019) for each donor. Donors’ expression maps were z-scored and winsorized (threshold: absolute z-score = 3.5). Co-expression was calculated separately for each gene and functional area. mRNA co-expression was calculated as the z-transformed Pearson correlation between the area and each of the 132 target ROIs. Subsequently, we calculated the Pearson correlation between co-expression and mean FC for each target ROI and averaged this across network. For each network and neurotransmitter, we tested whether co-expression-FC correlations differed from zero using one-sample t-tests. For the networks showing non-zero correlations of gene co-expression with FC, differences in the correlation between mRNA co-expression and functional connectivity were compared using Kolmogorov-Smirnov tests for each gene’s associated neurotransmitter (GABA or Glu). For the vAt specifically, we tested whether coupling between canonical structural and individual functional aspects was lower for Glu compared to GABA. To this end, we calculated bootstrapped mean correlations of gene co-expression associations with metabolite levels and mean correlations of FC with metabolite levels (number of iterations: 50,000). We resampled the resulting bootstrapped vectors and calculated the distance between functional and canonical associations, for both glutamate and GABA separately. Statistical significance of the difference between glutamate and GABA was assessed with a one-sample t-test.

#### Cognitive state correlates of p32 and p24 FC-glutamate association

To explore the potential functional relevance of the differential prediction of pgACC glutamate levels, we correlated the PLSR beta maps produced by models ran on functional *p24* and *p32*, as well as the difference between the two beta maps with association *Z* maps, using the Neurosynth decoder (see section *Methods and Materials, Neurosynth decoding*).

## Acknowledgements

We thank Renate Blober-Lüer and Dr. Claus Tempelmann (Department of Neurology, OVGU) for MR data acquisition. We would also like to thank Larissa Katz for her help with the analysis of MRS data.

This research was supported by the German Research Foundation (SFB779/A06 and DFG Wa2673/4-1 to M.W.) the Center for Behavioral Brain Sciences (CBBS NN05 to M.W.) and the University of Tübingen, Faculty of Medicine (fortune grant #2453-0-0 to N.B.K. and V.T.).

## Author contributions

M.W. designed the study to collect the data. L.M. and N.B.K. conceived of the method, analyzed the data, and interpreted the results and M.W. contributed to the interpretation. V.T. reviewed the code used for preprocessing and statistical analyses of the data. M.L. preprocessed the MRS data. N.P.G. provided the cytoarchitectonic parcellation of the pgACC. L.M. drafted the first version of the manuscript and N.B.K. contributed to the draft. All other authors revised the manuscript for important intellectual content.

---

**Supplementary Figure 1.**
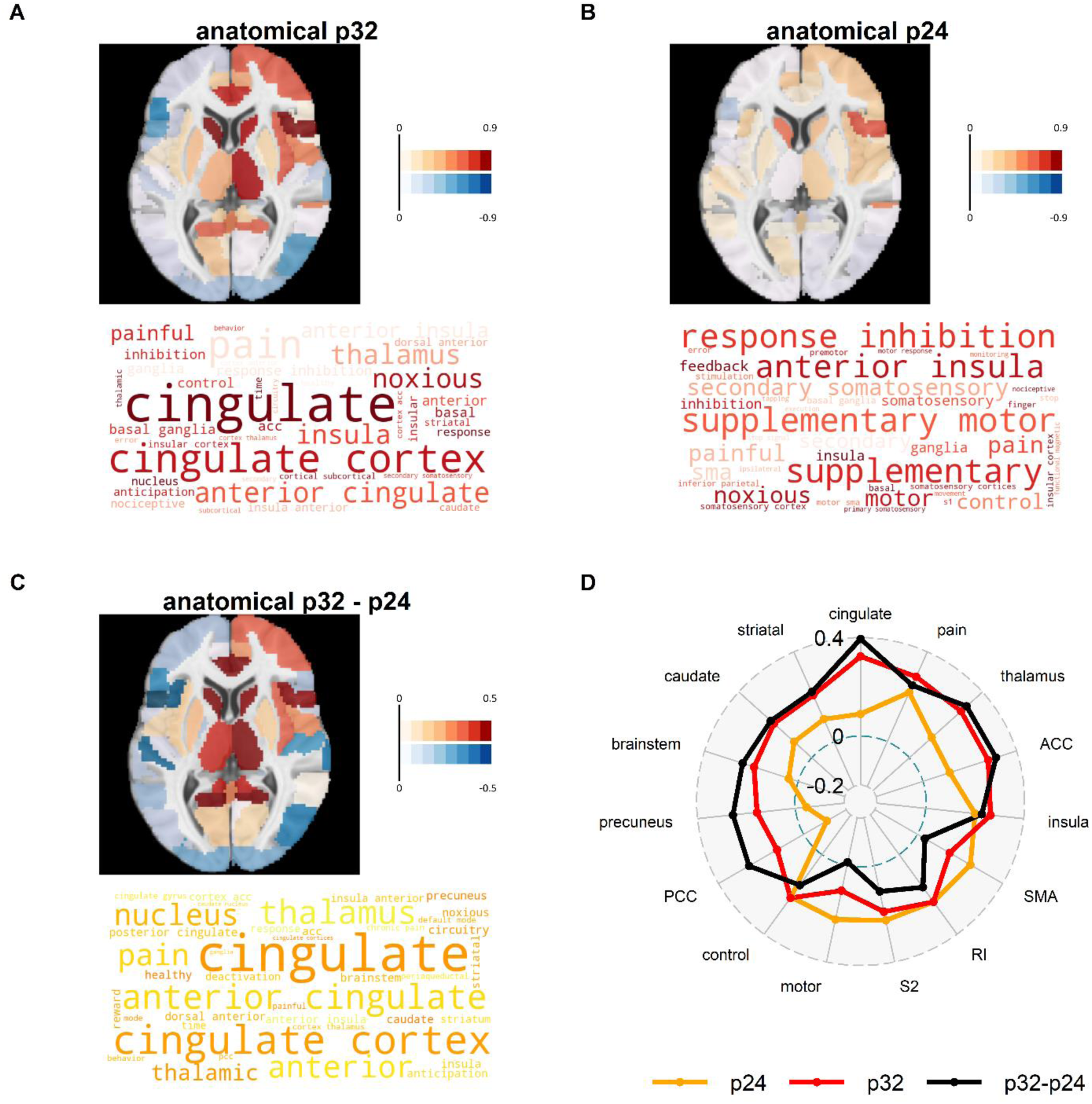
A-C, top: PLSR beta maps show the association between pgACC glutamate levels and pgACC area-to-target FC. A-C, bottom: decoded PLSR beta weight maps show the 40 cognitive states most strongly associated with the PLSR beta weight maps. D: spider plot of correlations with the top 5 terms (excluding duplicates) associated with PLSR beta weight maps of *p32*, *p24*, and *p32*-*p24* PLSR, respectively. ACC: “anterior cingulate”; SMA: “supplementary motor”; RI: “response inhibition”; S2: “secondary somatosensory”; PCC: “posterior cingulate”.

**Supplementary Figure 2.**
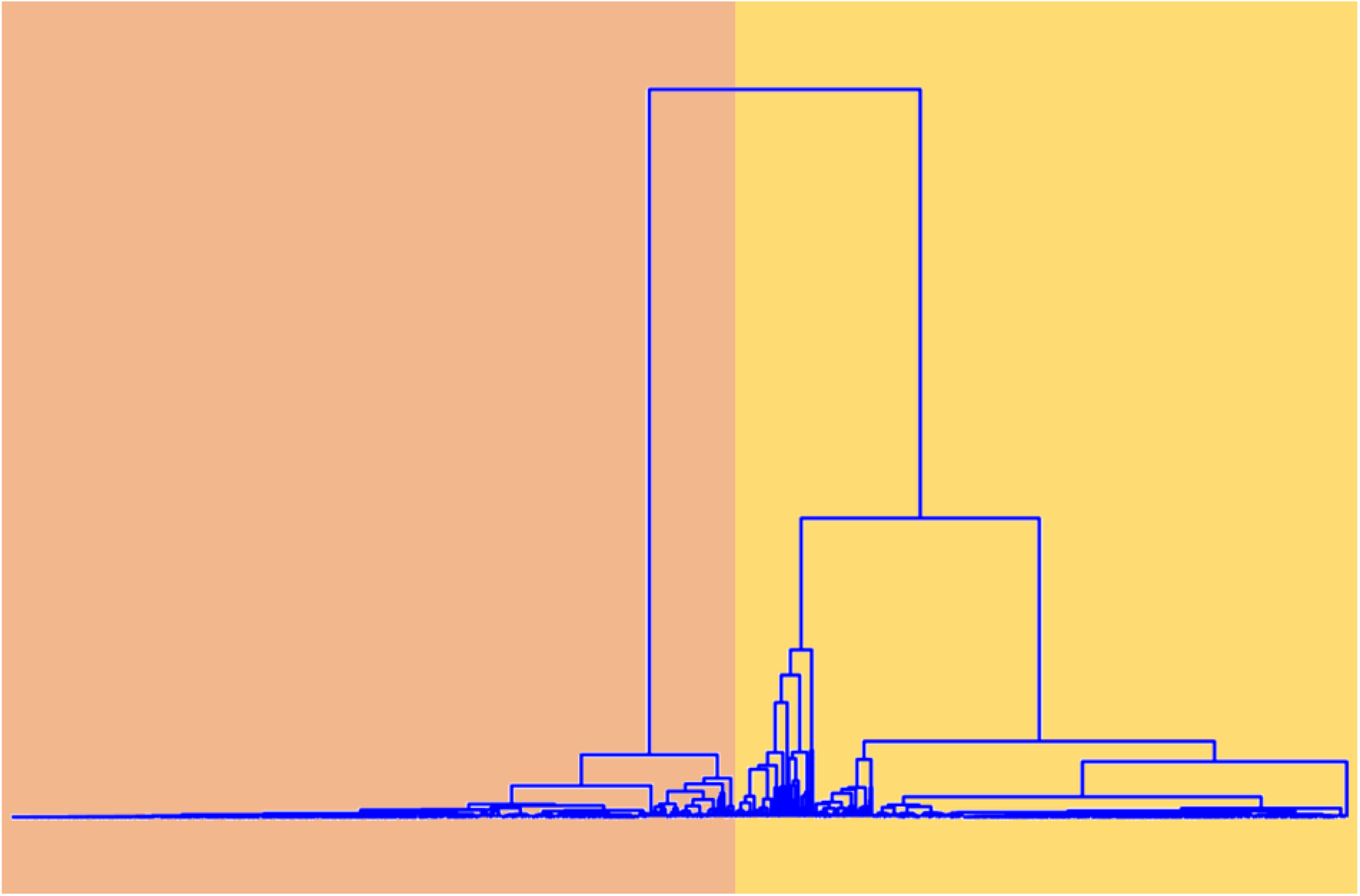
Dendrogram showing the results of hierarchical clustering of the MRS ROI based on functional connectivity to 132 target ROIs derived from the CONN atlas. Two main clusters emerged. Peach: functional *p32*; yellow: functional *p24*.

**Supplementary Table 1.**
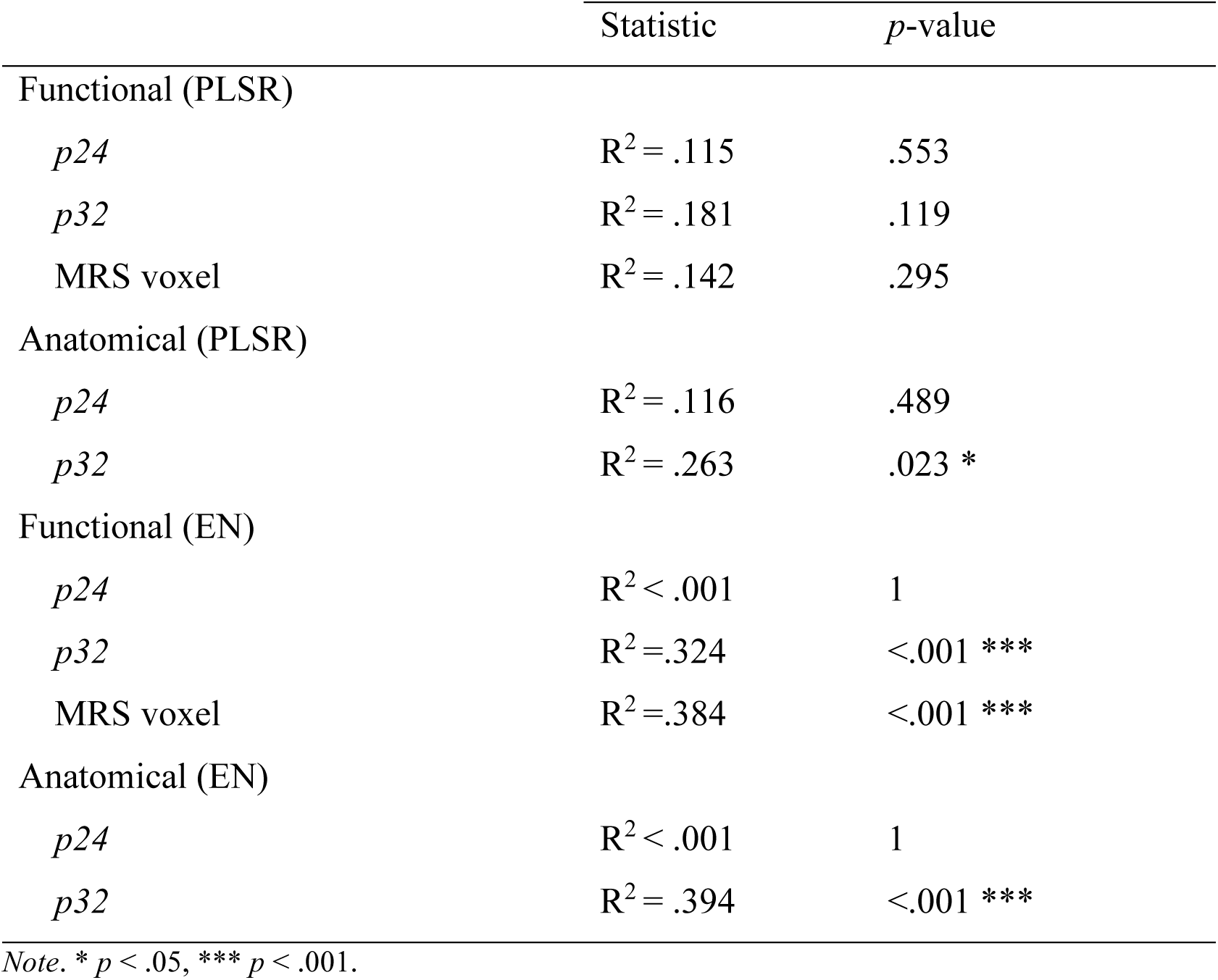
Prediction of Glu/tCr (residualized for GM %)

|  | Statistic | *p*-value |
| --- | --- | --- |
| Functional (PLSR) |  |  |
| *p*24 | $R^2 = .115$ | .553 |
| *p*32 | $R^2 = .181$ | .119 |
| MRS voxel | $R^2 = .142$ | .295 |
| Anatomical (PLSR) |  |  |
| *p*24 | $R^2 = .116$ | .489 |
| *p*32 | $R^2 = .263$ | .023 * |
| Functional (EN) |  |  |
| *p*24 | $R^2 < .001$ | 1 |
| *p*32 | $R^2 = .324$ | <.001 *** |
| MRS voxel | $R^2 = .384$ | <.001 *** |
| Anatomical (EN) |  |  |
| *p*24 | $R^2 < .001$ | 1 |
| *p*32 | $R^2 = .394$ | <.001 *** |

Note. \* $p < .05$ , \*\*\* $p < .001$ .

**Supplementary Table 2.** Demographics and effects of age on the prediction of Glu/tCr

|  | Statistic | *p*-value |
| --- | --- | --- |
| Glu/tCr |  |  |
| Age | $r(86) = -.256$ | .016* |
| Gray matter % | $r(86) = .015$ | .889 |
| Gender | $t(71.393) = .367$ | .715 |
| GABA/tCr |  |  |
| Age | $r(81) = .026$ | .819 |
| Gray matter % | $r(81) = -.064$ | .564 |
| Gender | $t(72.540) = -.717$ | .475 |
| Age |  |  |
| PLSR – cluster p32 | $R^2 = .210$ | .066 |
| PLSR – cluster p24 | $R^2 = .143$ | .289 |
| EN – cluster p32 | $R^2 = .441$ | < .001*** |
| EN – cluster p24 | $R^2 = .301$ | < .001*** |
| Glu/tCr and FC residualized for age |  |  |
| PLSR – cluster p32 | $R^2 = .174$ | .123 |
| PLSR – cluster p24 | $R^2 = .189$ | .084 |
| EN – cluster p32 | $R^2 = .423$ | < .001*** |
| EN – cluster p24 | $R^2 = < .001$ | 1 |

*Note.* Predictions of age and Glu/tCr were done using the FC from functional areas *p24* and *p32*. The models in which the influence of age was regressed out, used the residuals of Glu/tCr after regressing out both voxel gray matter content and age. \* $p < .05$ , \*\*\* $p < .001$ .

**Supplementary Table 3.** Prediction of GABA/tCr (residualized for GM %)

|  | Statistic | *p*-value |
| --- | --- | --- |
| Functional (PLSR) |  |  |
| *p*24 | $R^2 = .185$ | .068 * |
| *p*32 | $R^2 = .093$ | .637 |
| MRS voxel | $R^2 = .101$ | .609 |
| Anatomical (PLSR) |  |  |
| *p*24 | $R^2 = .126$ | .409 |
| *p*32 | $R^2 = .115$ | .503 |
| Functional (EN) |  |  |
| *p*24 | $R^2 = < .001$ | 1 |
| *p*32 | $R^2 = < .001$ | 1 |
| MRS voxel | $R^2 = < .001$ | 1 |
| Anatomical (EN) |  |  |
| *p*24 | $R^2 = < .001$ | 1 |
| *p*32 | $R^2 = < .001$ | 1 |

Note. \* $p < .05$ , \*\* $p < .01$ .

**Supplementary table 4.** Selected genes from the Allan Human Brain Atlas

| Gene | Entrez Gene ID | AHBA Probe ID | Neurotransmitter system |
| --- | --- | --- | --- |
| ABAT | 18 | 1010530 | GABA |
| GABBR1 | 2550 | 1023290 | GABA |
| GABBR2 | 9568 | 1048425 | GABA |
| GABRA1 | 2554 | 1014606 | GABA |
| GABRA2 | 2555 | 1013901 | GABA |
| GABRA3 | 2556 | 1056789 | GABA |
| GABRA4 | 2557 | 1056703 | GABA |
| GABRA5 | 2558 | 1056626 | GABA |
| GABRA6 | 2559 | 1056615 | GABA |
| GABRB1 | 2560 | 1056611 | GABA |
| GABRB2 | 2561 | 1056610 | GABA |
| GABRB3 | 2562 | 1029788 | GABA |
| GABRD | 2563 | 1056608 | GABA |
| GABRE | 2564 | 1025161 | GABA |
| GABRG1 | 2565 | 1029994 | GABA |
| GABRG2 | 2566 | 1027605 | GABA |
| GABRG3 | 2567 | 1021254 | GABA |
| GABRQ | 55879 | 1041466 | GABA |
| GAD1 | 2571 | 1056577 | GABA |
| GAD2 | 2572 | 1013593 | GABA |
| SLC32A1 | 140679 | 1034964 | GABA |
| SLC6A1 | 6529 | 1031772 | GABA |
| SLC6A11 | 6538 | 1051443 | GABA |
| SLC6A13 | 6540 | 1051441 | GABA |
| ALDH4A1 | 8659 | 1028289 | Glutamate |
| GLS | 2744 | 1056381 | Glutamate |
| GLS2 | 27165 | 1044598 | Glutamate |
| GLUD2 | 2747 | 1056377 | Glutamate |
| GLUL | 2752 | 1024398 | Glutamate |
| GOT1 | 2805 | 1056299 | Glutamate |
| GOT2 | 2806 | 1056295 | Glutamate |
| GRIA1 | 2890 | 1056189 | Glutamate |
| GRIA2 | 2891 | 1012842 | Glutamate |
| GRIA3 | 2892 | 1056188 | Glutamate |
| GRIA4 | 2893 | 1013883 | Glutamate |
| GRID1 | 2894 | 1028948 | Glutamate |
| GRID2 | 2895 | 1056185 | Glutamate |
| GRIK1 | 2897 | 1026823 | Glutamate |
| GRIK2 | 2898 | 1056180 | Glutamate |
| GRIK3 | 2899 | 1056123 | Glutamate |
| GRIK4 | 2900 | 1055987 | Glutamate |
| GRIK5 | 2901 | 1055952 | Glutamate |
| GRIN1 | 2902 | 1055939 | Glutamate |
| GRIN2A | 2903 | 1010806 | Glutamate |
| GRIN2B | 2904 | 1055935 | Glutamate |
| GRIN2C | 2905 | 1024347 | Glutamate |
| GRIN2D | 2906 | 1055933 | Glutamate |
| GRIN3A | 116443 | 1036035 | Glutamate |
| GRIN3B | 116444 | 1025776 | Glutamate |
| GRINA | 2907 | 1023366 | Glutamate |
| GRM1 | 2911 | 1010994 | Glutamate |
| GRM2 | 2912 | 1023800 | Glutamate |
| GRM3 | 2913 | 1055924 | Glutamate |
| GRM4 | 2914 | 1055922 | Glutamate |
| GRM5 | 2915 | 1015143 | Glutamate |
| GRM6 | 2916 | 1021370 | Glutamate |
| GRM7 | 2917 | 1025003 | Glutamate |
| GRM8 | 2918 | 1018070 | Glutamate |
| SLC17A7 | 57030 | 1040965 | Glutamate |
| SLC17A8 | 246213 | 1010718 | Glutamate |
| SLC1A1 | 6505 | 1025608 | Glutamate |
| SLC1A2 | 6506 | 1051582 | Glutamate |
| SLC1A3 | 6507 | 1051579 | Glutamate |
| SLC1A4 | 6509 | 1014665 | Glutamate |
| SLC1A5 | 6510 | 1019400 | Glutamate |
| SLC1A6 | 6511 | 1051575 | Glutamate |
| SLC1A7 | 6512 | 1051572 | Glutamate |

*Note:* AHBA = Allan Human Brain Atlas.

**Supplementary Table 5.** One sample t-tests for mRNA co-expression/FC correlations

|  | Glutamate-associated FC/co-expression |  |  | GABA-associated FC/co-expression |  |  |
| --- | --- | --- | --- | --- | --- | --- |
|  | *T*(85) | *p* | 95% *CI* | *T*(47) | *p* | 95% *CI* |
| Vis | -1.737 | .086 | [-0.113; 0.008] | -1.220 | .228 | [-0.118; 0.029] |
| SoM | 1.268 | .208 | [-0.017; 0.078] | 0.832 | .410 | [-0.042; 0.102] |
| dAt | 1.900 | .061 | [-0.004; 0.177] | 4.053 | <.001*** | [0.119; 0.354] |
| vAt | 21.916 | <.001*** | [0.506; 0.608] | 19.094 | <.001*** | [0.577; 0.713] |
| Lim | 1.064 | .290 | [-0.027; 0.089] | 2.839 | .007** | [0.026; 0.155] |
| FPN | 6.902 | <.001*** | [0.210; 0.380] | 5.810 | <.001*** | [0.220; 0.453] |
| DMN | 17.657 | <.001*** | [0.298; 0.373] | 13.850 | <.001*** | [0.365; 0.489] |
| Cer | 1.453 | .150 | [-0.013; 0.083] | 0.506 | .615 | [-0.047; 0.080] |

*Note.* Results of one-sample t-tests testing whether mRNA co-expression correlations with FC are different from zero, for each network and neurometabolite. Vis = visual network, SoM = somatomotor network, dAt = dorsal attention network, vAt = ventral attention network; Lim = limbic network; FPN = frontoparietal network; DMN = default mode network; Cer = cerebellar network. \*\* $p < .05$ , \*\*\* $p < .001$ .

**Supplementary Table 6.** GABA vs Glu: mRNA co-expression-FC correlations

|  | *p*24 |  | *p*32 |  | *p*24 + *p*32 |  |
| --- | --- | --- | --- | --- | --- | --- |
|  | D | *p* | D | *p* | D | *p* |
| vAt | 0.267 | .186 | 0.411 | .007** | 0.283 | .011* |
| FPN | 0.198 | .536 | 0.157 | .808 | 0.147 | .488 |
| DMN | 0.360 | .027* | 0.402 | .009** | 0.360 | <.001*** |

*Note.* Two-sample Kolmogorov-Smirnov tests comparing the distributions of z-transformed GABAergic and glutamatergic mRNA-co-expression and FC correlations. Vis = visual network, SoM = somatomotor network, dAt = dorsal attention network, vAt = ventral attention network; Lim = limbic network; FPN = frontoparietal network; DMN = default mode network; Cer = cerebellar network. \* $p < .05$ , \*\* $p < .01$ , \*\*\* $p < .001$ .

**Supplementary Table 7.** One-sample t-tests for PLSR beta weights (p24 + p32)

| Network<br>(df) | Glutamate |  |  | GABA |  |  |
| --- | --- | --- | --- | --- | --- | --- |
|  | *T* | *p* | 95% CI | *T* | *p* | 95% CI |
| vAt (13) | 4.839 | <.001*** | [0.209; 0.545] | 2.771 | .016 * | [0.046; 0.375] |
| FPN (13) | 3.910 | .002 ** | [0.118; 0.411] | 1.314 | .212 | [-0.077; 0.317] |
| DMN (39) | -0.356 | .724 | [-0.083; 0.058] | 0.482 | .633 | [-0.075; 0.122] |

*Note.* Results of one-sample t-tests testing whether PLSR beta weights (pooled across *p24* and *p32*) are different from zero, for each network and neurometabolite. vAt = ventral attention network; FPN = frontoparietal network; DMN = default mode network. \* $p < .05$ , \*\* $p < .01$ , \*\*\* $p < .001$ .

